# The developing brain structural and functional connectome fingerprint

**DOI:** 10.1101/2021.03.08.434357

**Authors:** Judit Ciarrusta, Daan Christiaens, Sean P. Fitzgibbon, Ralica Dimitrova, Jana Hutter, Emer Hughes, Eugene Duff, Anthony N Price, Lucilio Cordero-Grande, J-Donald Tournier, Daniel Rueckert, Joseph V Hajnal, Tomoki Arichi, Grainne McAlonan, A David Edwards, Dafnis Batalle

**Affiliations:** Department of Forensic and Neurodevelopmental Science, Institute of Psychiatry, Psychology and Neuroscience, King’s College London, London, United Kingdom; Centre for the Developing Brain, School of Imaging Sciences & Biomedical Engineering, King’s College London, London, United Kingdom; Department of Electrical Engineering, ESAT/PSI, KU Leuven, Leuven, Belgium; Wellcome Centre for Integrative Neuroimaging, FMRIB, Nuffield Department of Clinical Neurosciences, University of Oxford, UK; Paediatric Neuroimaging Group, Department of Paediatrics, University of Oxford, UK; Biomedical Image Technologies, ETSI Telecomunicación, Universidad Politécnica de Madrid & CIBER-BBN, Madrid, Spain; Biomedical Image Analysis Group, Department of Computing, Imperial College London, London, United Kingdom; Klinikum Rechts der Isar, Technical University of Munich, Munich, Germany; Department of Bioengineering, Imperial College London, London, SW7 2AZ, United Kingdom; Children’s Neurosciences, Evelina London Children’s Hospital, Guy’s and St Thomas’ NHS Trust, London, United Kingdom; MRC Centre for Neurodevelopmental Disorders, King’s College London, London, United Kingdom

## Abstract

In the mature brain, structural and functional connectivity ‘fingerprints’ can be used to identify the uniqueness of an individual. However, whether the characteristics that make a brain distinguishable from others already exist at birth remains unknown. Here, we used neuroimaging data of preterm born neonates who were scanned twice during the perinatal period to assess the developing brain fingerprint. We found that 62% of the participants could be identified based on the congruence of the later structural connectome to the initial connectivity matrix derived from the earlier timepoint. In contrast, similarity between functional connectomes only allowed to identify 12% of the participants. These suggests that structural connectivity is more stable in early life and can represent a potential connectome fingerprint. Thus, a relatively stable structural connectome appears to support a changing functional connectome at a time when neonates must rapidly acquire new skills to adapt to their new environment.

## INTRODUCTION

Advances in neuroimaging technology have provided new means to investigate the human brain *in vivo*. This has enabled characterization of a connectome delineating the structural and functional organization of the brain at a macro-scale. This connectome can be represented as a large-scale correlation matrix where each row and column correspond to brain subunit indices which may be structural or functional nodes, so that each matrix element describes ‘connectivity’ between two neural parts (Sporns, Tononi and Kötter, 2005; Sporns, 2010). The information contained in the functional and structural connectome of an individual is highly specific to that person and has been compared to a personal ‘fingerprint’ (Finn *et al.*, 2015; Yeh *et al.*, 2016). Although, the functional connectome has been demonstrated to be highly stable over multiple years after late adolescence (Horien *et al.*, 2019), a delay in establishing a distinctive functional connectome through adolescence has been linked to mental health difficulties (Kaufmann *et al.*, 2017). However, the structural and functional connectome of an infant differs from older age groups (Cao, Huang and He, 2017), and the extent to which either is stable (i.e., reproducible at the level of the individual) is unknown. A better understanding of the extent of malleability of a given property of an individual’s brain and its relation to outcomes may guide personalized approaches to optimize child neurodevelopmental health.

The developing brain is governed by dynamic processes that transform an amalgam of a few cells in early gestation into a complex organ capable of rapidly processing and integrating information. The foetal period is marked by cortical migratory processes, with anomalies at this stage frequently leading to neuronal migration disorders and atypical brain structure (Ten Donkelaar and Van der Vliet, 2004; Kostović *et al.*, 2014). A hallmark of the third foetal trimester is a change in the relative proportion of short and long range association fibres (Ouyang *et al.*, 2019). These fast-changing microscopic developmental mechanisms quickly lead to reshaping of macrostructural features, which can be captured with Magnetic Resonance Imaging (MRI). For example, despite ongoing rapid growth, cortical folding features show remarkable high similarity between scans of the same subject across the first year following birth, suggesting a high self-similarity in cortical macrostructure between birth and the first 2 postnatal years (Duan et al., 2020). However, the stability of structural/functional network metrics and whether the connectome is ‘individual’ in early life has not been previously investigated. This is important to understand because, although the structural and functional connectome are inter-related and complimentary, they represent models of brain connectivity with distinct developmental influences (for a review see (Suárez *et al.*, 2020)).

Useful information about the perinatal brain and its subsequent maturation has already been acquired. In the perinatal brain, in parallel with structural changes, cortical neurons fire specific activity patterns which further regulate genetic expression and refine structure into functional systems (Tau and Peterson, 2010; Kirkby *et al.*, 2013). Highly connected structural and functional nodes/hubs are crucial for information flow and further evolve as myelination matures (Morgan *et al.*, 2018). A striking difference between the neonatal and the adult brain is that functional hubs, highly interconnected regions, are more restricted to somatosensory, auditory, visual and motor regions, unlike the higher order networks seen in adults (Cao *et al.*, 2017). In childhood, the identification accuracy of the functional connectome at rest has been estimated to be at 43% (Vanderwal *et al.*, 2019), relative to 92% reported in adults (Finn *et al.*, 2015), suggesting acquiring functional diversity and consequent uniqueness is part of the trajectory towards adulthood. However, the structural connectome fingerprint has only been investigated in adults and been shown to be relatively plastic globally but very stable within specific white matter bundles, such as the corpus callosum (Yeh *et al.*, 2016). On the contrary, the neonatal brain is still immature and differs from the adult one, so it is possible that nature and extent of identifiable features of individuals’ structural and functional connectomes that are stable over time also differ.

Here, we investigate whether a structural and/or functional fingerprint is already established perinatally, by assessing the similarity of the structural and functional connectome of preterm born infants, who were scanned soon after birth and then again at term equivalent age. Unlike adults, the maturing brain is highly dynamic and undergoing rapid reorganization. Therefore, we hypothesized that connectome similarity would be lowest when the time between scans was longest. In addition, brain activity becomes experience-driven during this early postnatal period and experiences increase daily as the infant interacts with the world ex-utero (Greenough, Black and Wallace, 1987; Khazipov and Luhmann, 2006). Thus, we also hypothesized that this constantly changing functional activity leads to experience-dependent changes in the functional connectome upon a (relatively) stable structural connectome.

## METHODS

### Subjects

Research participants were prospectively recruited as part of the developing Human Connectome Project (dHCP), an observational, cross-sectional Open Science programme approved by the UK National Research Ethics Authority (14/LO/1169). Written consent was obtained from all participating families prior to imaging.

As part of the dHCP, a total of 63 subjects were scanned twice and had both functional and diffusion MRI data acquired. After pre-processing, 18 diffusion datasets were discarded due to poor registration to template space, resulting in a total of 45 subjects (26 males) with good quality diffusion MRI data. The infants were born at a median of 32.29 weeks gestational age (GA) [range: 25.57-37], with their first scan acquired at a median of 35 weeks post menstrual age (PMA) [range: 29.29-37.43] and the second acquired at a median of 41 weeks PMA [range: 38.43-44.86]. From the functional data, 18 subjects were discarded due to poor registration to template space and additional 14 had to be discarded due to significant signal loss during acquisition. Thus, there was a total of 31 subjects (21 males) with good quality functional connectome data, born at a median of 34.57 weeks GA [range: 27.57-37.00], had their first scan at a median of 35.57 weeks PMA [range: 30.86-37.43] and had their second scan at a median of 40.57 weeks PMA [range: 38.86-44.57]. The final sub-group with both good quality structural and functional data consisted of 26 subjects (16 males), born at a median age of 34.14 weeks [range: 28.71-37.00], with their first scan acquired at a median of 35.43 weeks PMA [range: 31.43-37.43] and the second acquired at a median of 40.93 weeks PMA [range: 38.86-44.86] (details described in Table 1).

**Table 1.**
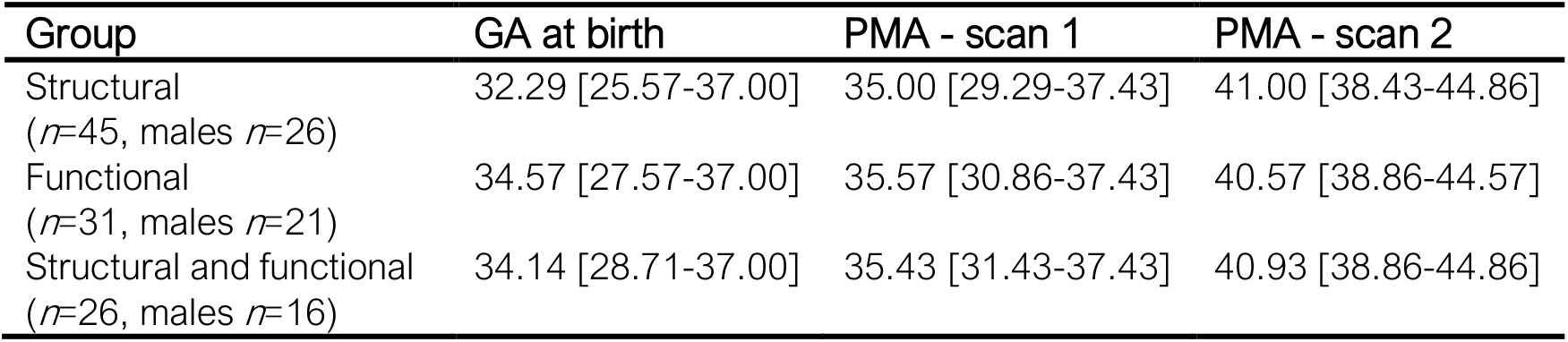
Descriptive sample characteristics

### Data acquisition

All subjects underwent Magnetic Resonance Imaging (MRI) scanning at the Evelina Newborn Imaging Centre, St Thomas’ Hospital, London, UK. Structural, diffusion and functional data was acquired using a 3 Tesla Philips Achieva system (Philips Medical Systems, Best, The Netherlands) with customized neonatal imaging system including a 32-channel phased-array head coil (Rapid Biomedical, Rimpar, Germany) (Hughes *et al.*, 2017). Infants were studied during natural sleep following feeding and immobilization in a vacuum evacuated bag (Med-Vac, CFI Medical Solutions, Fenton, MI, USA). Hearing protection (moulded dental putty in the external auditory meatus (President Putty, Coltene Whaledent, Mahwah, NJ, USA) and earmuffs (MiniMuffs, Natus Medical Inc., San Carlos, CA, USA)) and physiological monitoring (oxygen saturations, heart rate, axillary temperature) were applied before data acquisition. MR-compatible foam shielding was used for further acoustic noise attenuation. All scans were supervised by a neonatal nurse and/or paediatrician who monitored heart rate, oxygen saturation and temperature throughout the scan.

A total of 300 volumes of diffusion MRI were acquired over 19 minutes and 20 seconds with b-values of 400 s/mm^2^, 1000 s/mm^2^ and 2600 s/mm^2^ spherically distributed in 64, 88 and 128 directions respectively, with 20 b=0 s/mm^2^ images and parameters: Multiband factor 4, SENSE factor 1.2, partial Fourier 0.86, acquired in-plane resolution 1.5×1.5mm, 3mm slices with 1.5mm overlap, repetition time (TR)/ echo time (TE) of 3800/90ms and 4 phase-encoding directions (Hutter *et al.*, 2018). After reconstruction, image resolution was 1.5 mm isotropic. High-temporal-resolution BOLD fMRI optimized for neonates was acquired over 15 minutes 3 seconds (2300 volumes) using a multislice gradient-echo echo planar imaging (EPI) sequence with multiband excitation (multiband factor 9), TR 392 ms, TE 38 ms, flip angle 34°, and acquired spatial resolution 2.15 mm isotropic (Price *et al.*, 2015). For clinical interpretation and registration purposes, a Turbo spin echo sequence (parameters: TR = 12s, TE = 156ms, SENSE factor 2.11 (axial) and 2.54 (sagittal)) was used to acquire high-resolution T2-weighted (T2w) images. The T2w axial and sagittal volumes originally acquired at 0.8×0.8mm, 1.6mm slices with 0.8mm overlap were motion corrected and super-resolved to a final resolution of 0.8mm isotropic (Cordero-Grande *et al.*, 2018).

### Image pre-processing and connectome construction

A neonatal specific segmentation pipeline (Makropoulos *et al.*, 2014) was used to obtain tissue segmentation of each subject’s T2w images in native space. A neonatal adaptation (Shi *et al.*, 2011) of the AAL atlas (Tzourio-Mazoyer *et al.*, 2002) aligned to the dHCP high-resolution neonatal template (Schuh *et al.*, 2018) was used to parcellate each’s subject brain into 90 cortical and subcortical regions. Previously calculated tissue segmentation and T2w images were used as input for a non-linear registration based on a diffeomorphic symmetric image normalization method (SyN) available in ANTS software (Avants *et al.*, 2011) to bring the 90 regions neonatal atlas into the subject’s native space (Supplementary Table 1).

Pre-processing of diffusion MRI data and structural connectome construction was performed as previously reported (Taoudi-Benchekroun *et al.*, 2020). Briefly, after hybrid SENSE reconstruction (Zhu *et al.*, 2016), diffusion signal was denoised (Cordero-Grande *et al.*, 2019), and susceptibility distortions were corrected (Andersson, Skare and Ashburner, 2003). A spherical harmonics and radial decomposition (SHARD) slice-to-volume reconstruction was applied to further correct motion effects and other artefacts (Christiaens *et al.*, 2021). The N4 algorithm (Tustison *et al.*, 2010) implemented in MRtrix (Tournier *et al.*, 2019) was applied for bias field correction. Multi-tissue CSD (Jeurissen *et al.*, 2014) using restricted anisotropic diffusion for brain tissue and free diffusion for fluid like features (Pietsch *et al.*, 2019) was used to estimate fibre orientation distribution (FOD) in each brain voxel. Response functions for each tissue type were generated as the average from the response functions in an independent sub-group of 20 healthy term control neonates from the dHCP (Taoudi-Benchekroun *et al.*, 2020). Multi-tissue log-domain intensity normalisation (Raffelt, et al. 2017) was applied to FODs, and normalised brain tissue like FODs were used to generate 10 million streamlines with anatomically constrained probabilistic tractography (Smith et al., 2012) with biologically accurate weights (SIFT2) (Smith, et al. 2015). The fibre density SIFT2 proportionality coefficient (μ) for each subject was obtained to achieve inter-subject connection density normalisation. Atlas parcellation and tissue maps in T2w native space were registered to diffusion space with a rigid registration using b=0 volumes as target. (Schnabel *et al.*, 2001). The structural connectome of each infant was constructed in native diffusion space, by calculating the μ × SIFT2-weighted sum of streamlines connecting each pair of regions into a weighted adjacency matrix of size 90×90.

For functional data, all 2300 volumes of fMRI data acquired per participant were utilized without undergoing any scrubbing. Data were pre-processed using the Developing Human Connectome Project pipeline optimized for neonatal fMRI, detailed in (Fitzgibbon *et al.*, 2020). In brief, susceptibility dynamic distortion together with intra- and inter-volume motion effects were corrected in each subject using a bespoke pipeline including slice-to-volume and rigid-body registration (Andersson *et al.*, 2017). In order to regress out signal artifacts related to head motion, cardiorespiratory fluctuations and multiband acquisition, 24 extended rigid-body motion parameters were regressed together with single-subject ICA noise bespoke components identified with the FSL FIX tool (Oxford Centre for Functional Magnetic Resonance Imaging of the Brain’s Software Library, version 5.0) (Salimi-Khorshidi *et al.*, 2014). Atlas parcellation and tissue maps were propagated from T2w native space using a boundary-based registration (Greve and Fischl, 2009). Average timeseries of each of the previously parcelled ROIs intersecting with GM regions were calculated in native fMRI space. Functional connectivity (FC) was calculated as the Pearson’s partial correlation of the signal between each pair of ROIs, controlling for the signal in the rest of the ROIs (Shi et al., 2011) and resulting in a matrix of size 90×90. Negative correlations were not considered and set to zero.

### Similarity Analysis

#### Global similarity - Identifiability rate quantification

To identify the optimal network density threshold for structural and functional connectome similarity, we tested the percentage of correctly identified subjects for all possible thresholds (Supplementary Figure 1). Based on this analysis, we applied a 25% network density threshold to the structural and functional connectomes at time-point 1 for each subject. This connectivity matrix was then binarized and used as a mask for the connectivity matrix in time-point 2 ensuring that the same inter-regional connections were compared between the two time points. Spearman’s correlation between each matrix at time-point 1 and time-point 2 was calculated, resulting in a similarity matrix of 45×45 subjects for structural connectivity and 31×31 subjects for functional connectivity. If the self-similarity between time-point 1 and time-point 2 (diagonal correlation) was higher than the self-to-other-similarity, this was quantified as a successful match (Finn *et al.*, 2015).

For visualization purposes, all similarity values were normalised by scaling relative to the maximum correlation between time-point 1 and all other subjects at time-point 2 dividing by the maximum value in each row (i.e., for each row, value of 1 indicated the maximum match between time-point 1 and time-point 2). As shown in Figure 1 this scaling results in a value of 1 in the diagonal when self-similarity is higher than any self-to-other-similarity value. If the value of 1 is not in the diagonal it indicates self-to-other-similarity is higher than self-similarity.

**Figure 1.**
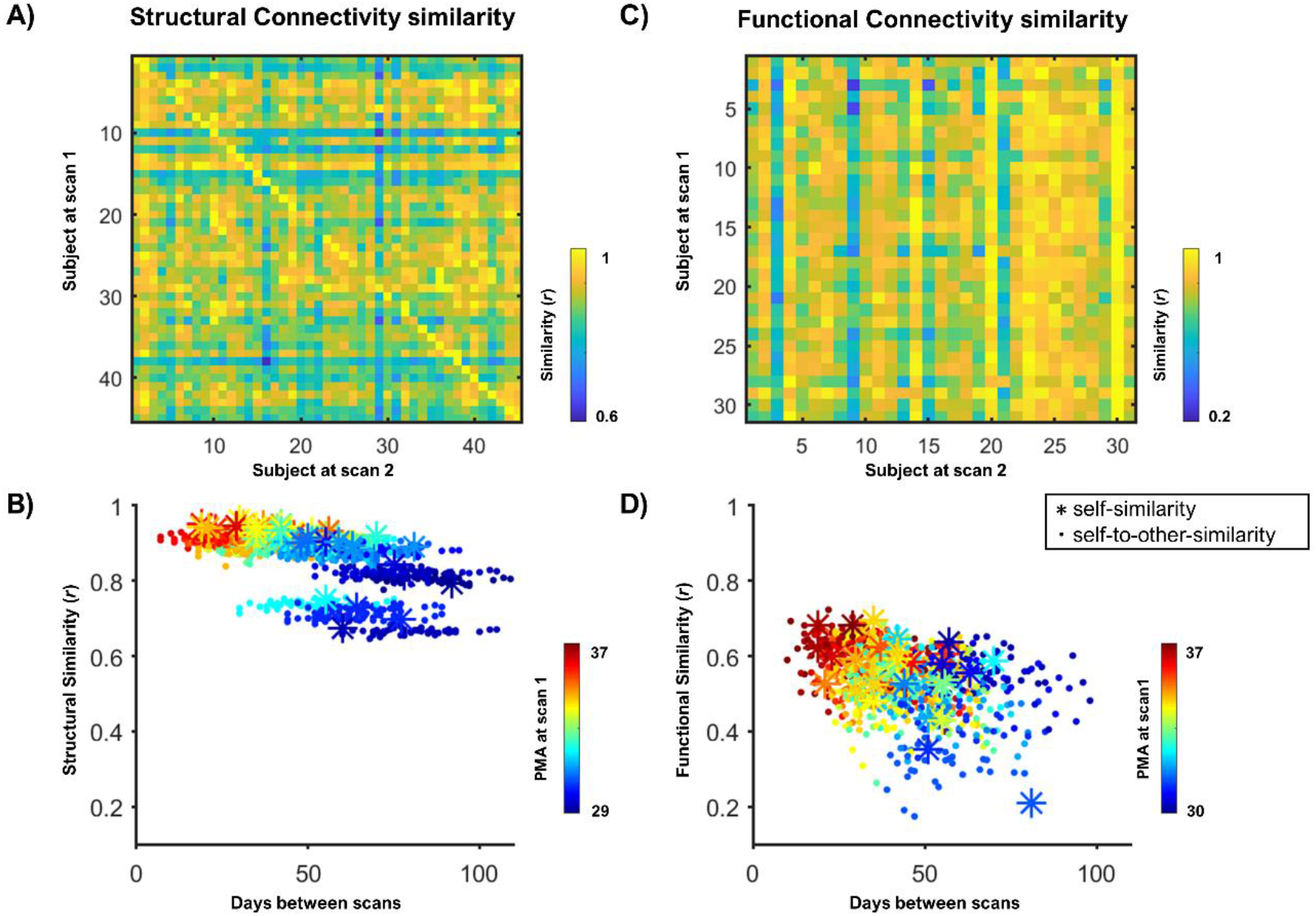
Structural and Functional global similarity. The correlation between the connectome of each subject at time-point 1 and 2 is depicted by the similarity matrix for structural connectivity (A) and functional connectivity (C). The correlations are then plotted against days between scans with a colour gradient showing the age of the subject at time-point 1 for structural data (B) and functional data (D). The stars represent the correlation between the connectome of a subject at time-point 1 with the connectome of the same subject at time-point 2 (i.e., self-similarity), and the dots represent the correlation of a subject at time-point 1 with a different subject at time-point 2 (i.e., self-to-other-similarity).

The same analysis was repeated for a sub-group of participants that had both structural and functional connectome data. This resulted in two 26×26 matrices containing structural and functional data, characterising the identifiability rate for the two modalities in the same individuals. The diagonal values of these matrices were extracted to provide median and range of structural and functional self-similarity.

#### Age effect on self-similarity and self-to-other-similarity

In order to assess the effect of age in the structural and functional fingerprint, we first calculated the partial correlation between PMA at scan at time-point 1 and self-similarity (controlling for days between scans), and then the partial correlation between days between scans and self-similarity (controlling for PMA at time-point 1) for the whole group in each modality. Then, for the sub-group we converted the self-similarity and the self-to-other-similarity into z-scores for better visualization of the effect of age on self-similarity. If the self-similarity z-score of a subject was higher than any of the self-to-other-similarity z-score, this would be equivalent to successfully matching a subject between time-point 1 and time-point 2, as in previous fingerprinting studies (Finn *et al.*, 2015).

To further characterize the effect of age on self-similarity, we ran a general linear model analysis with PMA at time-point 1 and days between scans as independent variables and self-similarity as dependent variable. This allowed quantification of the beta coefficients of the age effect on self-similarity. To characterize self-to-other-similarity association with age difference between time-point 1 and time-point 2, we performed a linear mixed-effects model (LME) with days between scans as fixed effect and with subject at time-point1 dependent random effect for the intercept to account for repeated measures.

#### Regional analysis

Given the distinct trajectories of maturation of subcortical and cortical regions, we repeated all the analysis described above in 7 clusters that represent larger anatomical areas: central, frontal, limbic, occipital, parietal, deep grey matter and temporal. These regions were composed of 8, 22, 14, 14, 10, 8, and 14 nodes respectively. The central cluster was composed of bilateral precentral and postcentral gyrus, paracentral lobules and supplementary motor areas. The frontal cluster was composed of superior, middle, inferior and orbitofrontal cortices, as well as the olfactory lobule and the rectus gyrus. The limbic cluster was composed of the insula, anterior and posterior cingulate, hippocampus and amygdala. The occipital cluster was composed of the calcarine, cuneus, lingual cortices together with superior, inferior and middle occipital gyrus and the fusiform. The parietal cluster was composed of the superior and inferior parietal gyrus, supramarginal, angular and precuneus gyrus. The deep grey matter cluster was composed of all basal ganglia structures and thalami. The temporal cluster was composed of the Rolandic operculum, the Heschl gyrus, and superior, middle and interior temporal cortices as well as the temporal poles. Supplementary Table 1 contains all 90 regions of the atlas and to which cluster they belong to. This allowed to calculate similarity identifiability rates, similarity z-scores and age linear models for each sub-region.

### Data availability

The dHCP is an open-access project. The imaging and collateral data can be downloaded by registering at https://data.developingconnectome.org/.

Derived data including structural and functional connectivity networks used in this study are available in github.com/code-neuro/neonatalconnectomefingerprint/.

## RESULTS

### Whole-brain similarity

The structural connectome comparison between the scans at preterm and term equivalent ages yielded strong correlations not only among the scans of the same subject but also between the scans of different subjects (Figure 1AB). The mean self-similarity was *r*=0.90, ranging from *r*=0.67 to *r*=0.96, and the mean self-to-other-similarity was *r*=0.88, ranging from *r*=0.65 to *r*=0.94. The identifiability rate was 28/45 (62.22%), representing the percentage of times self-similarity was higher than the self-to-other-similarity. Furthermore, we found a significant correlation between PMA at first scan and self-similarity values (i.e., the older a subject was at the first scan, the more “self-similar” their structural connectome was between scans), controlling for days between scans (*r*=0.49, *p*=0.006).

The functional connectome comparison showed less consistent results (Figure 1CD). The correlation values between scans were lower compared to structural connectome similarities. The mean self-similarity was *r*=0.56, ranging from *r*=0.21 to *r*=0.70, and the mean self-to-other-similarity was *r*=0.54, ranging from *r*=0.17 to *r*=0.73. Thus, the identifiability rate for functional connectome similarity was 3/31 (9.68%). We observed no significant correlation between PMA at first scan and functional self- similarity, but there was a significant negative correlation between functional self-similarity and days between scans (*r*=-0.39, *p*=0.03).

Closer inspection of the self-similarity for the structural and functional connectivity in a sub-group of participants with data available in both modalities confirmed that all subjects had higher similarity between their structural connectomes than their functional connectomes. The median structural self-similarity was *r*=0.93, ranging from *r*=0.89 to *r*=0.96, and the median functional self-similarity was *r*=0.55, ranging from *r*=0.21 to *r*=0.69. The identifiability rate for structural data was 18/26 (69.23%), in contrast with 3/26 (11.54%) for functional data (Figure 2). As the structural and functional data examined were from the same individuals, age and time between scans was exactly matched.

**Figure 2.**
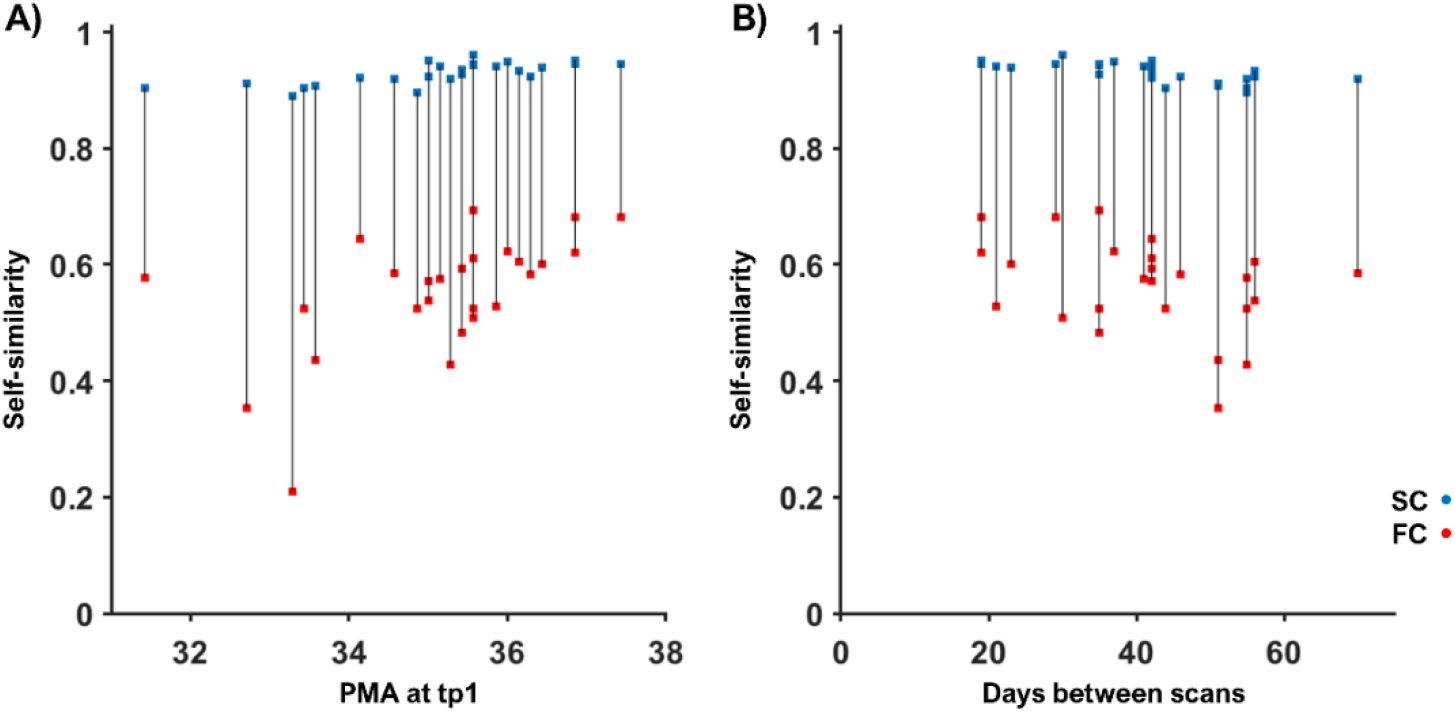
Self-similarity for structural and functional connectivity. The similarity correlation between the structural connectivity (SC) matrices between scans (blue) and between the functional connectivity (FC) matrices (red) is plotted again age at first scan (A) and against days between scans (B).

Finally, we converted the self-similarity and self-to-other-similarity values into z-scores for each subject and sorted them based on age at time-point 1 to better visualize whether older subjects have a more identifiable whole-brain structural connectome (Figure 3). Results show that the structural connectome was more stable than the functional connectome against variations on age or time between scans.

**Figure 3.**
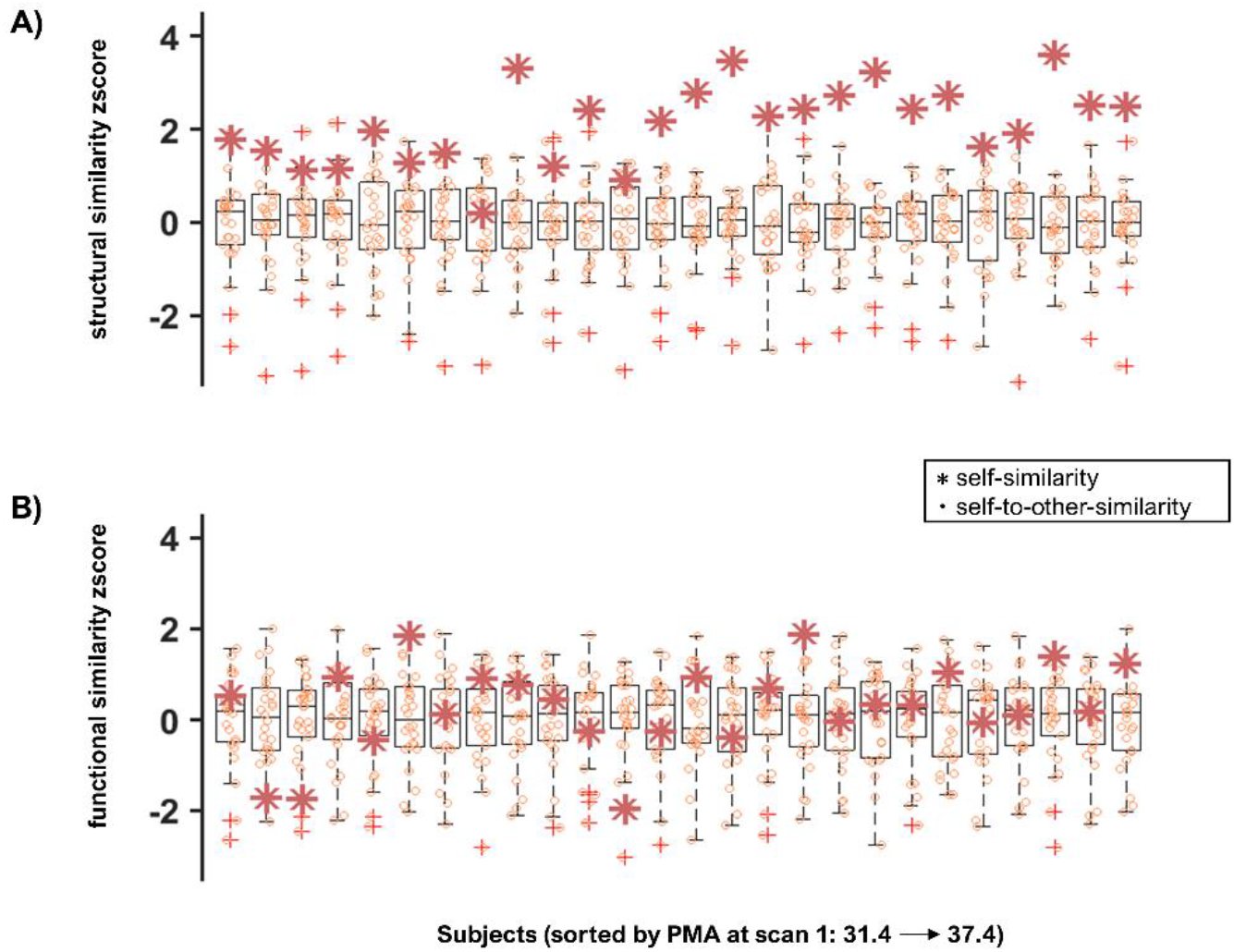
Self-similarity and self-to-other-similarity z-scores arranged by PMA. The boxplots show the similarity scores between time-point 1 and time-point 2 converted to z-scores for each participant arranged from left to right (youngest to oldest at time-point 1). The stars represent self-similarity and the circles represent self-to-other-similarity. The upper row depicts structural connectome similarity (A) and the bottom row shows functional connectome similarity (B).

### Age effect on sub-group similarity

The general linear model analysis for the sub-group with both structural and functional data further showed that age at time-point 1 has a significant effect on global structural connectome self-similarity (β=0.007, p=0.01). We observed no significant effect of days between scans on structural connectome self-similarity; and no effect of age at time point 1 or days between scans on global functional connectome self-similarity (Table 2).

**Table 2.** Beta coefficients and p-values for the effect of age and days between scans on self-similarity.

|  | *SC* |  |  |  | *FC* |  |  |  |
| --- | --- | --- | --- | --- | --- | --- | --- | --- |
|  | PMA |  | Days between scan |  | PMA |  | Days between scan |  |
| | $\beta$ | p | $\beta$ | p | $\beta$ | p | $\beta$ | p |
| *global* | 0.00611 | 0.01 | -0.00058 | 0.01 | 0.02413 | 0.15 | -0.00265 | 0.08 |
| *central* | 0.00126 | 0.72 | -0.00024 | 0.44 | -0.03211 | 0.06 | -0.00258 | 0.09 |
| *frontal* | 0.00031 | 0.91 | -0.00044 | 0.07 | 0.02662 | 0.31 | -0.00270 | 0.25 |
| *limbic* | 0.00444 | 0.28 | -0.00069 | 0.07 | 0.01875 | 0.28 | -0.00187 | 0.23 |
| *occipital* | -0.00107 | 0.74 | -0.00046 | 0.13 | 0.06104 | 0.01 | -0.00007 | 0.97 |
| *parietal* | -0.00002 | 1.00 | 0.00029 | 0.60 | 0.08789 | 0.00 | 0.00174 | 0.47 |
| *sub-cortical* | -0.00150 | 0.61 | -0.00019 | 0.47 | -0.00186 | 0.91 | -0.00069 | 0.64 |
| *temporal* | 0.00216 | 0.44 | -0.00015 | 0.56 | 0.01117 | 0.54 | -0.00041 | 0.80 |

The LME analysis showed age at first scan also has a significant effect on global structural connectome self-to-others-similarity (β_2_=0.006, p<0.001), but there was no significant effect of days between scans. Unlike what was observed for self-similarity, both days between scans (β_1_=-0.001, p<0.001) and age at time-point 1 (β_2_=0.02, p<0.001) had a significant effect on global functional connectome self-to-others-similarity.

### Regional similarity

Regional comparison of structural connectomes showed lower identifiability rate compared to the global metrics. The limbic cluster (insula, cingulate, hippocampus and amygdala) showed highest identifiability rate at 11/45 (24.44%), followed by frontal regions at 10/45 (22.22%), occipital at 7/45 (15.56%), central at 5/45 (11.11%), both parietal and deep GM at 4/45 (8.89%) and temporal at 3/45 (6.67%).

Functional connectome comparisons yielded similar identifiability rates compared to whole-brain metrics. They were still qualitatively lower than structural similarity identifiability rates, except for the parietal cluster. Central and parietal regions showed the highest identifiability rate of 3/31 (9.68%). Limbic regions showed an identifiability rate of 2/31 (6.45%) and frontal, occipital, sub-cortical and temporal clusters showed an identifiability rate of 1/31 (3.23%) (Figure 4).

**Figure 4.**
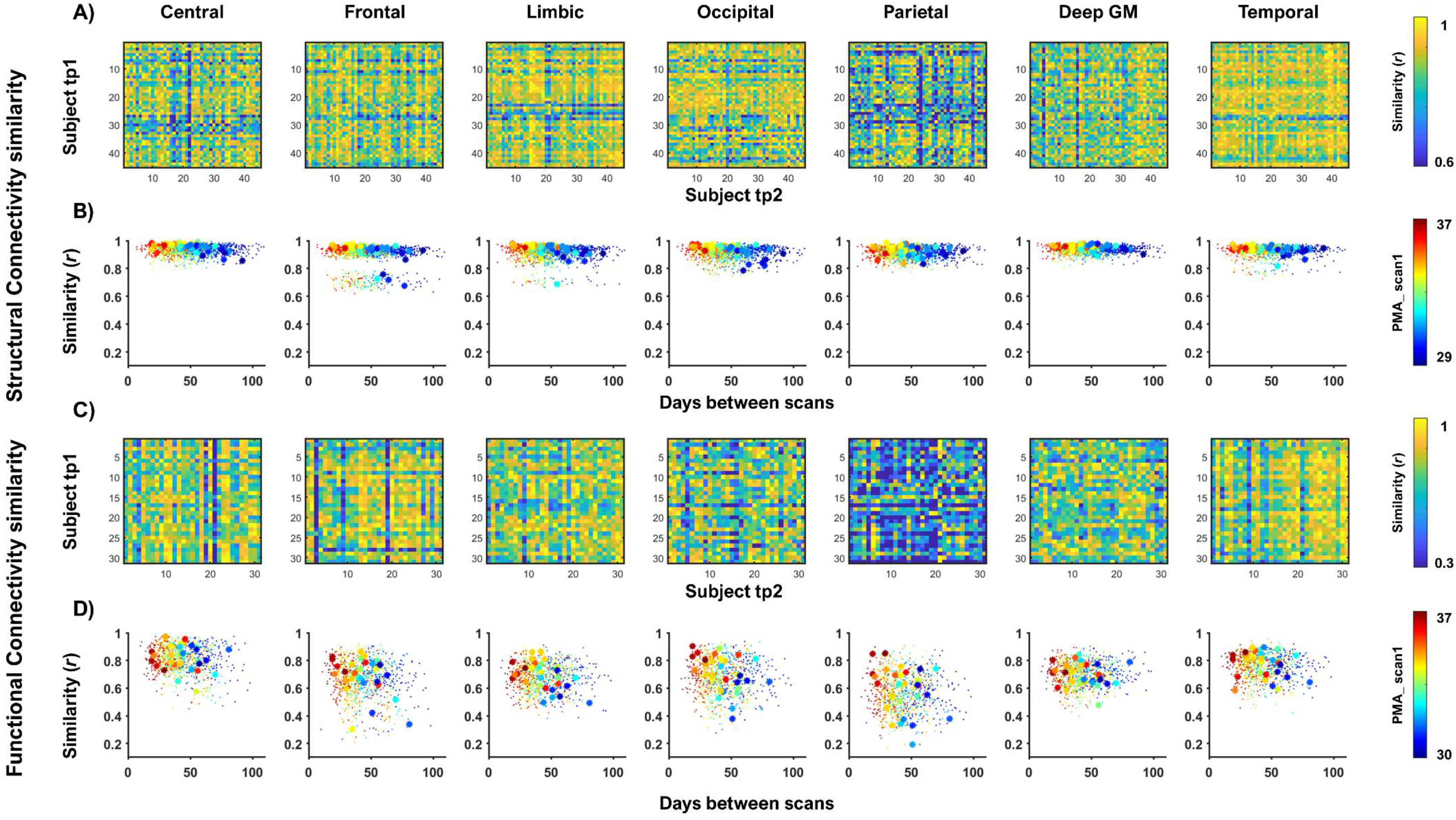
Structural and Functional cluster-wise similarity. Similarity matrices together with plots to depict the association of the similarity with days between scans underneath are shown for structural connectivity (A, B) and functional connectivity (C, D). These figures are presented in different columns for different anatomical clusters: somatosensory-motor or central region, frontal, limbic, occipital, parietal, deep grey matter, and temporal.

Closer qualitative inspection of the self-similarity for the structural and functional connectivity in the sub-group of participants with both structural and functional connectomes showed structural and functional self-similarity closer together in the central cluster and more dispersed in the frontal cluster (Figure 5). However, functional similarity identifiability values were always lower than structural identifiability rates in this sub-group.

**Figure 5.**
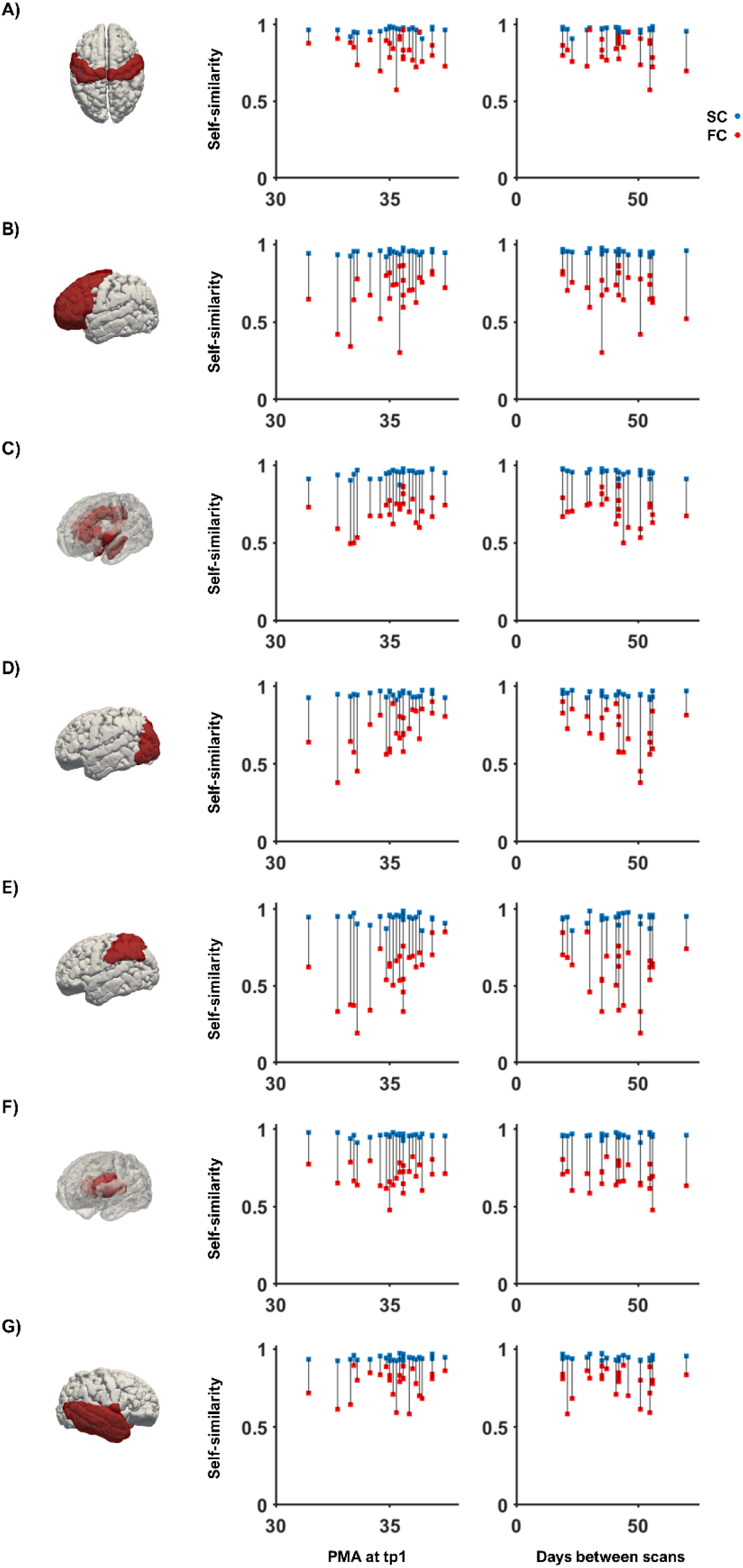
Regional self-similarity for structural and functional connectivity cluster-wise. The structural self-similarity (blue) and the functional self-similarity (red) is plotted against age at first scan (PMA at time-point 1) in the first column and against days between scans in the second column for central (A), frontal (B), limbic (C), occipital (D), parietal (E), deep grey matter (F) and temporal (G) cluster. Grey lines provide visual guidance to match structural and functional similarity values of the same subject.

### Age effect on regional self-similarity

The generalized linear model run independently for each cluster in the sub-group with both structural and functional data showed that age at time-point 1 has a significant effect on functional connectome similarity in the parietal region after Bonferroni correction (β=0.088, p=0.003). We observed no significant effect of age or days between scans on any other cluster for functional nor structural self-similarity after multiple comparisons correction (Table 2).

## DISCUSSION

In the current study we used a unique set of longitudinal high quality structural and functional neonatal brain MRI data from the Developing Human Connectome Project to investigate the status of the connectome fingerprint at an early stage of neurodevelopment. To do so, we selected data from preterm born infants that were scanned soon after birth and then again at term equivalent age. Our results show that the whole-brain structural connectome can already identify an older individual at birth. In contrast, the whole-brain functional connectome changes more noticeably between scan timepoints, so individual identification is less stable based on their functional connectome regardless of their age at birth or time between scans.

### A structural connectivity fingerprint is present during the perinatal period

During the perinatal period, the brain undergoes marked micro and macrostructural changes (Kunz *et al.*, 2014; Batalle *et al.*, 2017; Pietsch *et al.*, 2019).Nevertheless, our observations suggest that by the normal time of birth, an individual’s brain structural connectome is relatively stable. This suggests that the individual template of structural connectivity is predominately genetically determined, in the absence of an external insult. Consistent with this, macroscale structural white matter tractography has been shown to be highly heritable with axial diffusivity, radial diffusivity and fractional anisotropy of commissural fibres found to have the highest genetic influence and association fibres the least (Lee *et al.*, 2015). By term equivalent age, the neonatal brain has an established framework of thalamocortical fibres, a large abundance of u-shaped cortico-cortical fibres, and visible long range association pathways such as the cingulum bundle and callosal fibres (Takahashi *et al.*, 2012). The abundance of u-shaped cortico-cortical fibres at term is in line with an early establishment of cortical folding patterns that remain individually unique throughout the first two postnatal years (Duan *et al.*, 2020). Thus, the acquisition of a stable structural connectome appears to coincide with the attainment of more mature brain structural appearance and may therefore be a marker of maturity. This fits with the observed relationship between higher structural connectome self-similarity and an older age at the time of the first scan. Alongside self-similarity, we also observed high self-to-other-similarity values in the structural connectome which likely represents the development of common macroscale features (structural connectivity backbone).

### Functional connectivity fingerprint is absent at birth

Task-based fMRI studies in neonates have demonstrated that the primary sensory cortices (e.g. somatosensory, auditory, olfactory and visual) are capable of processing external stimuli and are undergoing activity-dependent maturation during the perinatal period (Anderson *et al.*, 2001; Arichi *et al.*, 2010; Perani *et al.*, 2010; Lee *et al.*, 2012; Baldoli *et al.*, 2015; Allievi *et al.*, 2016; Adam-darque *et al.*, 2018; Dall’Orso *et al.*, 2018). Resting state studies have reported significant age-dependent increases in functional short-range connectivity in the somatosensory, visual, auditory and language networks throughout the perinatal time window, further suggesting rapid functional plasticity and maturation within these systems (Cao *et al.*, 2017). These changes in local degree centrality may represent functional reshaping of primary resting state networks that resemble adult networks by term equivalent age, while higher-order association networks appear immature (Eyre *et al.*, 2020). Such fast reorganization across multiple functional systems during the perinatal time window may explain why we did not observe a single case where self-similarity was higher than self-to-others-similarity.

Our results markedly contrast with those of previously reported work in children. To our knowledge, the youngest previously investigated population for connectome similarity analysis to date has been a sample of 6-year-old children, where they reported an identification rate of 43% between resting state functional connectomes (Vanderwal *et al.*, 2019). Another study in children aged 7 to 15 years reported high correlation coefficients between functional connectivity matrices of the same participant, almost on par with adult similarity values (Horien *et al.*, 2019). While our results indicate that a functional connectome fingerprint is barely present at birth, future studies should investigate when this starts to increase between infancy and childhood.

### Impact of age at first scan and days between scans on similarity

To disentangle the impact of inter-scan interval and age at first scan on similarity of the structural and functional connectome, we examined a sub-group of 26 neonates who had both data types. We observed a significant effect of age at time-point 1 on the global structural self-similarity. This effect might be determined by the individual-specific developmental trajectory of white matter microstructure and not necessarily indicative of an adult-like unique structural connectome (Ouyang *et al.*, 2019).

The absence of any significant effect of age on global functional self-similarity at this early stage of development might be explained by the reshaping of long-range functional connectivity, which matures within the first postnatal year (Damaraju *et al.*, 2014). However, we cannot exclude the possibility that the low functional connectome self-similarity in the perinatal period is a false negative. For example, there may be a non-linear effect that cannot be captured with a linear model or an anatomical parcellation might not be optimal to characterize the stability of the functional connectome in early development. Future studies focusing in functional connectome similarity should investigate if an appropriate parcellation derived from functional data or multi-modal information based parcellations (Glasser *et al.*, 2016) yields higher self-similarity rates.

Interestingly, we observed the same age effect on global structural connectome self-to-other-similarity. This suggests certain age dependent organization patterns are also common across subjects. The same was observed for functional self-to-other-similarity, which significantly decreased with longer intervals between scans being compared and increased with older age at time point 1. The low self-similarity, combined with strong self-similarity-to others and its age dependency roots for time constrained functional changes at a developmental time where there is a strong mixture of finely tuned spontaneous neural activity patterns and sensory experience dependent mechanisms (Hanganu-Opatz, 2010; Toyoizumi *et al.*, 2013). The strong dependency of age and similarity patterns across subjects may also be related to genetic expression patterns that are highly dynamic and fast changing during this time of development (Silbereis *et al.*, 2015).

### Regional fingerprinting of the connectome

Brain development broadly follows a posterior to anterior maturational trajectory (Huttenlocher and Dabholkar, 1997) with sensory systems developing before higher order networks (Cao, Huang and He, 2017). Structural connectome identifiability rate was lower when investigating regions separately, suggesting global metrics with more data points is more informative as a fingerprint when using structural data. Later developing networks such as frontal and limbic cortices had the highest structural self-similarity. Given the late maturation of these regions, we speculate the high self-similarity might be driven by the fact that these regions are undergoing the least amount of structural change during the interval between scans.

We saw the highest identifiability rate in the central cluster of the functional connectome. This suggests that functional sensory-motor networks can provide higher identifiability rates in babies, while in comparison frontal-parietal structures appear more unique in adults (Finn *et al.*, 2015). However, it will require a larger sample of data specifically comparing somatosensory and frontal regions in infants and adults to test this hypothesis. In addition, structural and functional coupling is a sign of maturation which appears more robust after adolescence (Baum *et al.*, 2020). Thus, the closer resemblance between structure and function self-similarity in the central cluster might be indicative of the relatively mature state of somatosensory and motor cortices in the perinatal time window.

### Limitations

The main limitation of this study is that to assess the connectome fingerprint across weeks in the perinatal period *ex-utero* there is no other option but to investigate a preterm born population. Hence, to what extent the fingerprint is affected by preterm birth or is representative of normal development will have to be investigated in older infant cohorts or using foetal MRI. Another limitation pertains the multiple developmental factors which may influence the acquired signal in different ways and consequently the connectome. Despite the robustness of SIFT to characterize structural connections (Smith *et al.*, 2015), using the most advanced pipelines available for neonatal fMRI data processing (Fitzgibbon *et al.*, 2020), and being very stringent on data quality measures, the potential influence of developmental factors on the signal is out of our control. For diffusion data, developmental changes such as cortical folding or tissue water content reduction among others can affect the signal differently at different ages. Similarly, developmental effects which may influence the BOLD signal in different ways might relate to vascular density or neurovascular coupling, which can affect the sensitivity and specificity at different ages. Therefore, MR signal changes on the similarity values reported might be beyond differences in the neural “fingerprint”. On this line, it is also important to note we used anatomical parcellations to characterize the functional nodes and future studies should investigate whether diverse functional parcellation methods mimic or differ from the findings reported in this study.

## CONCLUSION

The brain structural connectome fingerprint is already present in the perinatal period. It is relatively stable and individually unique at this stage of development, while functional connectivity is either too dynamic or immature to provide strong identification features. Region-wise analysis suggested that the functional fingerprint in early development might be more stable within clusters, although identifiability rates were still higher for structural data. Future studies should investigate regional differences throughout development, the association of the global structural fingerprint to developmental outcome, and whether genetic or environmental risk impact the stability of the fingerprint.

## ACKNOWLEDGEMENTS

We would like to thank the infants and parents who contributed their time to this research. We are also thankful to our colleagues from the Newborn Imaging Centre at Evelina London. Conflict of Interest. None declared.

## Supplementary Figure

**Supplementary Figure 1.**
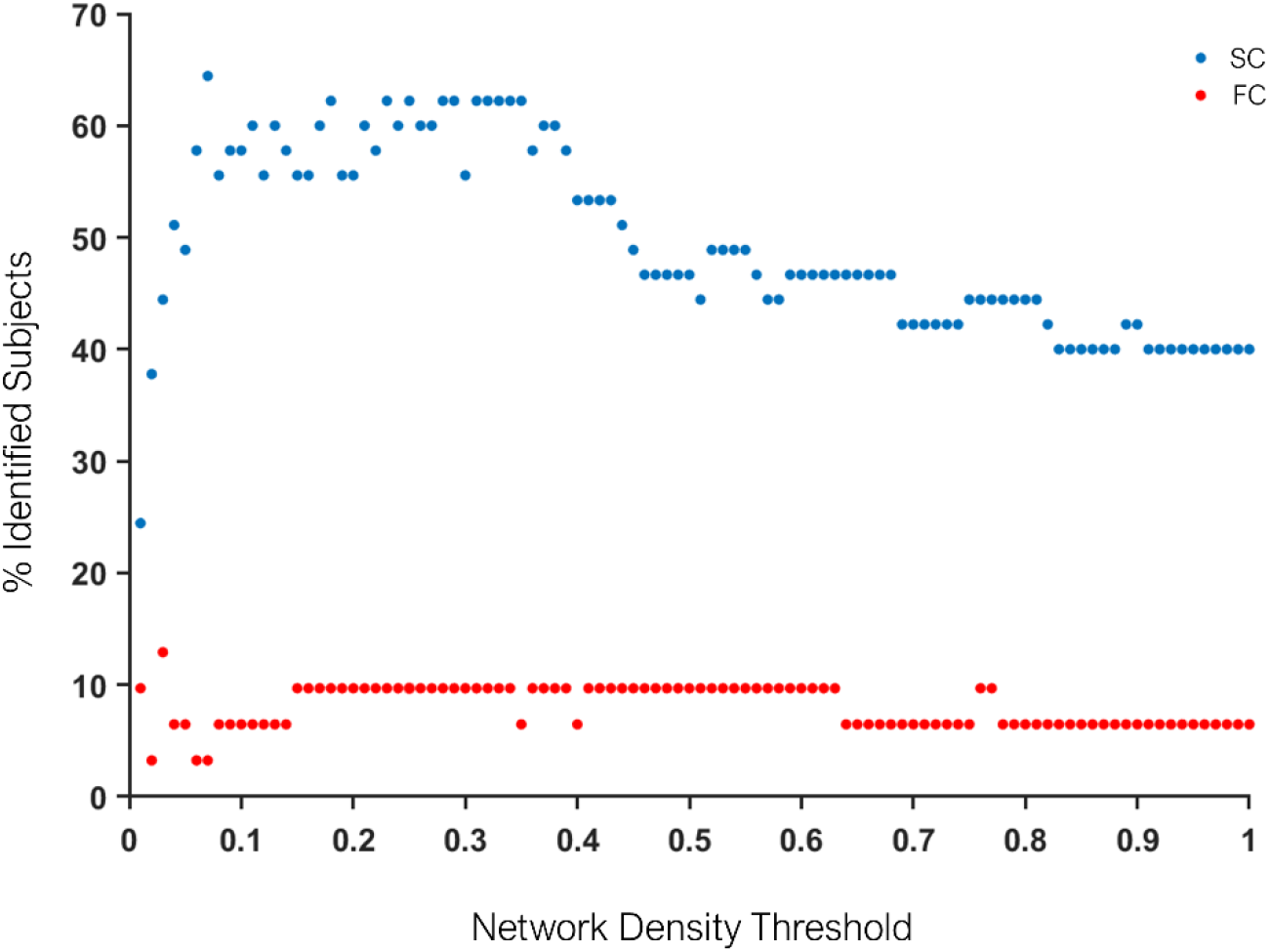
Network Density dependent identifiability rate for structural (blue - SC) and functional (red-FC) connectivity.

## Supplementary Table

**Supplementary Table 2.**
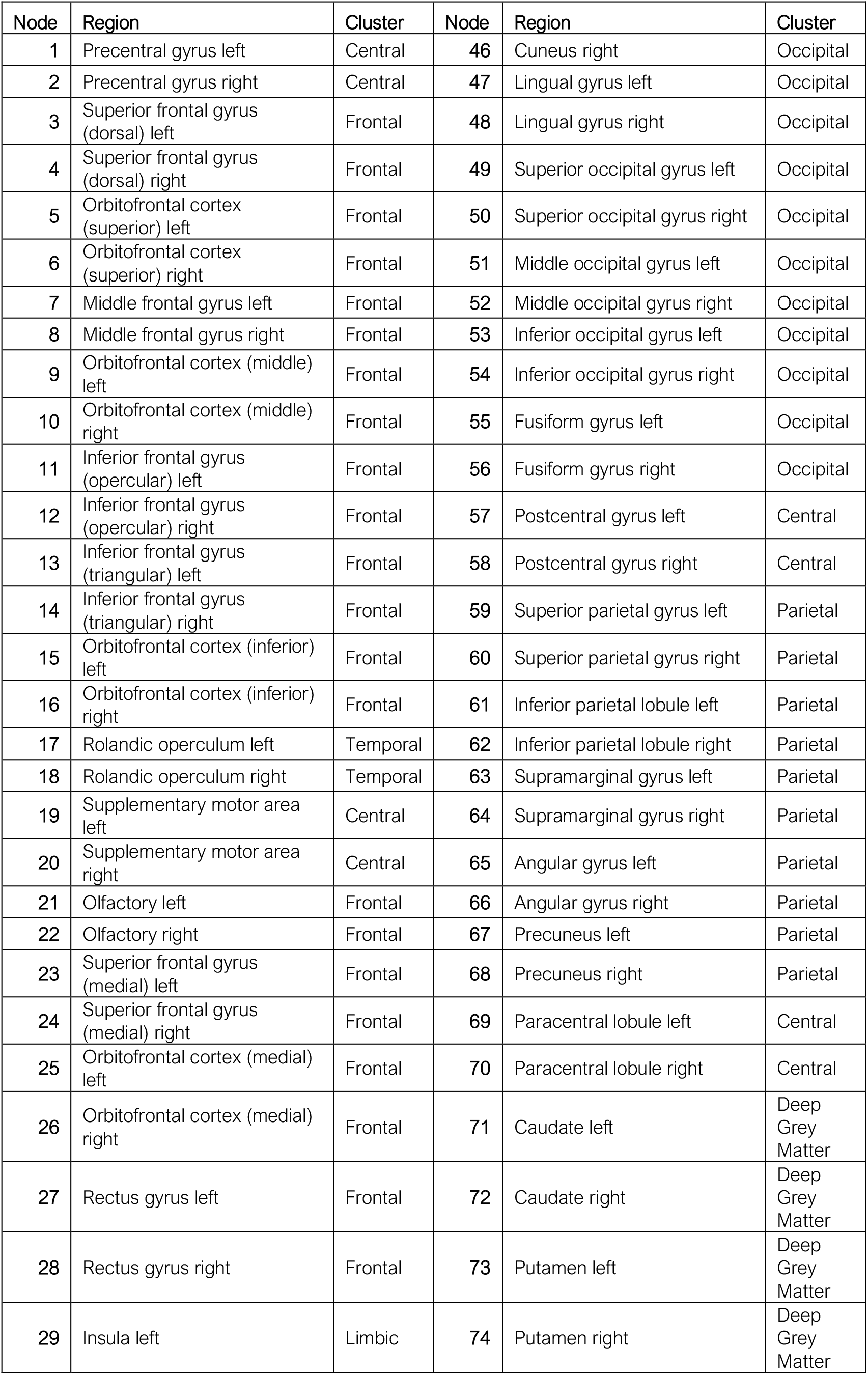
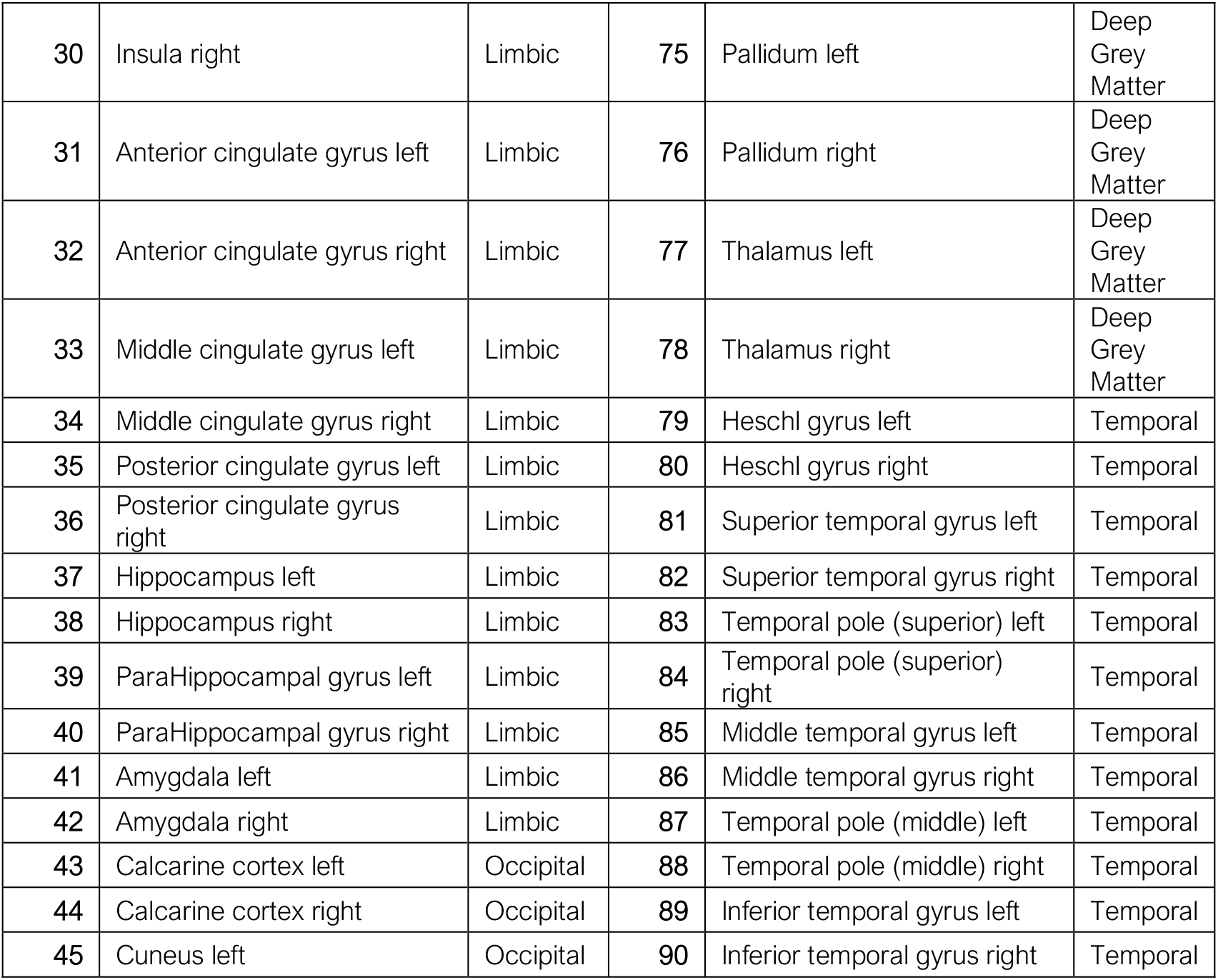
Connectome nodes: anatomical regions and clusters

## REFERENCES

Adam-darque, A. et al. (2018) ‘fMRI-based Neuronal Response to New Odorants in the Newborn Brain’, Cerebral Cortex, 28(8), pp. 2901–2907.

Allievi, A. G. et al. (2016) ‘Maturation of Sensori-Motor Functional Responses in the Preterm Brain’, Cerebral Cortex, 26(1), pp. 402–413.

Anderson, A. W. et al. (2001) ‘Neonatal auditory activation detected by functional magnetic resonance imaging’, Magnetic Resonance Imaging, 19(1), pp. 1–5.

Andersson, J. L. R. et al. (2017) ‘Towards a comprehensive framework for movement and distortion correction of diffusion MR images: Within volume movement’, NeuroImage. Elsevier, 152, pp. 450–466.

Andersson, J. L. R., Skare, S. and Ashburner, J. (2003) ‘How to correct susceptibility distortions in spin-echo echo-planar images: Application to diffusion tensor imaging’, NeuroImage, 20(2), pp. 870–888.

Arichi, T. et al. (2010) ‘Somatosensory cortical activation identified by functional MRI in preterm and term infants’, NeuroImage, 49(3), pp. 2063–2071.

Avants, B. B. et al. (2011) ‘A reproducible evaluation of ANTs similarity metric performance in brain image registration’, NeuroImage, 54(3), pp. 2033–2044.

Baldoli, C. et al. (2015) ‘Maturation of preterm newborn brains: a fMRI–DTI study of auditory processing of linguistic stimuli and white matter development’, Brain Structure and Function, 220(6), pp. 3733–3751.

Batalle, D. et al. (2017) ‘Early development of structural networks and the impact of prematurity on brain connectivity’, NeuroImage. Elsevier, 149, pp. 379–392.

Baum, G. L. et al. (2020) ‘Development of structure–function coupling in human brain networks during youth’, Proceedings of the National Academy of Sciences of the United States of America, 117(1), pp. 771–778.

Cao, M. et al. (2017) ‘Early Development of Functional Network Segregation Revealed by Connectomic Analysis of the Preterm Human Brain’, Cerebral cortex, 27(3), pp. 1949–1963.

Cao, M., Huang, H. and He, Y. (2017) ‘Developmental Connectomics from Infancy through Early Childhood’, Trends in Neurosciences. Elsevier Ltd, 40(8), pp. 494–506.

Christiaens, D. et al. (2021) ‘Scattered slice SHARD reconstruction for motion correction in multi-shell diffusion MRI’, NeuroImage. Elsevier Inc., 225, p. 117437.

Cordero-Grande, L. et al. (2018) ‘Three-dimensional motion corrected sensitivity encoding reconstruction for multi-shot multi-slice MRI: Application to neonatal brain imaging’, Magnetic Resonance in Medicine, 79(3), pp. 1365–1376.

Cordero-Grande, L. et al. (2019) ‘Complex diffusion-weighted image estimation via matrix recovery under general noise models’, NeuroImage.

Dall’Orso, S. et al. (2018) ‘Somatotopic Mapping of the Developing Sensorimotor Cortex in the Preterm Human Brain’, Cerebral Cortex, 28(7), pp. 2507–2515.

Damaraju, E. et al. (2014) ‘Functional connectivity in the developing brain: A longitudinal study from 4 to 9months of age’, NeuroImage. Elsevier Inc., 84, pp. 169–180.

Ten Donkelaar, H. J. and Van der Vliet, T. (2004) ‘Overview of the Development of the Human Brain and Spinal Cord’, in Clinical Neuroanatomy, pp. 1–45.

Duan, D. et al. (2020) ‘Individual identification and individual variability analysis based on cortical folding features in developing infant singletons and twins’, Human Brain Mapping, 41, pp. 1985–2003.

Eyre, M. et al. (2020) ‘The Developing Human Connectome Project: Typical and disrupted perinatal functional connectivity’, bioRxiv, (0).

Finn, E. S. et al. (2015) ‘Functional connectome fingerprinting : identifying individuals using patterns of brain connectivity’, Nature neuroscience. Nature Publishing Group, 18(11), pp. 1664–1671.

Fitzgibbon, S. P. et al. (2020) ‘The developing Human Connectome Project (dHCP) automated resting-state functional processing framework for newborn infants’, NeuroImage. Elsevier Inc., 223(August), p. 117303.

Glasser, M. F. et al. (2016) ‘A multi-modal parcellation of human cerebral cortex’, Nature. Nature Publishing Group, 536, pp. 171–178.

Greenough, W. T., Black, J. E. and Wallace, C. S. (1987) ‘Experience and Brain Development’,Child Development, 58, pp. 539–559.

Greve, D. N. and Fischl, B. (2009) ‘Accurate and robust brain image alignment using boundary-based registration’, NeuroImage. Elsevier Inc., 48(1), pp. 63–72.

Hanganu-Opatz, I. L. (2010) ‘Between molecules and experience: Role of early patterns of coordinated activity for the development of cortical maps and sensory abilities’, Brain Research Reviews, 64(1), pp. 160–176.

Horien, C. et al. (2019) ‘The individual functional connectome is unique and stable over months to years’, NeuroImage. The Authors, 189, pp. 676–687.

Hughes, E. J. et al. (2017) ‘A dedicated neonatal brain imaging system’, Magnetic Resonance in Medicine, 78, pp. 794–804.

Huttenlocher, P. R. and Dabholkar, A. S. (1997) ‘Regional differences in synaptogenesis in human cerebral cortex’, Journal of Comparative Neurology, 387(2), pp. 167–178.

Hutter, J. et al. (2018) ‘Time-Efficient and Flexible Design of Optimized Multishell HARDI Diffusion’, Magnetic Resonance in Medicine, 79, pp. 1276–1292.

Jeurissen, B. et al. (2014) ‘Multi-tissue constrained spherical deconvolution for improved analysis of multi-shell diffusion MRI data’, NeuroImage. Academic Press Inc., 103, pp. 411–426.

Kaufmann, T. et al. (2017) ‘Delayed stabilization and individualization in connectome development are related to psychiatric disorders’, Nature Neuroscience, 20(4), pp. 513–517.

Khazipov, R. and Luhmann, H. J. (2006) ‘Early patterns of electrical activity in the developing cerebral cortex of humans and rodents’, Trends in Neurosciences, 29(7), pp. 414–418.

Kirkby, L. A. et al. (2013) ‘A Role for Correlated Spontaneous Activity in the Assembly of Neural Circuits’, Neuron. Elsevier, 80(5), pp. 1129–1144.

Kostović, I. et al. (2014) ‘The Relevance of Human Fetal Subplate Zone for Developmental Neuropathology of Neuronal Migration Disorders and Cortical Dysplasia.’, CNS neuroscience & therapeutics, 21(3), pp. 1–9.

Kunz, N. et al. (2014) ‘Assessing white matter microstructure of the newborn with multi-shell diffusion MRI and biophysical compartment models’, NeuroImage. Elsevier B.V., 96, pp. 288–299.

Lee, S. J. et al. (2015) ‘Quantitative tract-based white matter heritability in twin neonates’, NeuroImage. Elsevier Inc., 111, pp. 123–135.

Lee, W. et al. (2012) ‘Visual functional magnetic resonance imaging of preterm infants’, Developmental Medicine and Child Neurology, 54(8), pp. 724–729.

Makropoulos, A. et al. (2014) ‘Automatic whole brain MRI segmentation of the developing neonatal brain’, IEEE Transactions on Medical Imaging. Institute of Electrical and Electronics Engineers Inc., 33(9), pp. 1818–1831.

Morgan, S. E. et al. (2018) ‘A Network Neuroscience Approach to Typical and Atypical Brain Development’, Biological Psychiatry: Cognitive Neuroscience and Neuroimaging. Elsevier Inc, 3(9), pp. 754–766.

Ouyang, M. et al. (2019) ‘Delineation of early brain development from fetuses to infants with diffusion MRI and beyond’, NeuroImage. Elsevier Ltd, 185, pp. 836–850.

Perani, D. et al. (2010) ‘Functional specializations for music processing in the human newborn brain’, Proceedings of the National Academy of Sciences, 107(10), pp. 4758–4763.

Pietsch, M. et al. (2019) ‘A framework for multi-component analysis of diffusion MRI data over the neonatal period’, NeuroImage. Elsevier Ltd, 186, pp. 321–337.

Price, A. et al. (2015) ‘Accelarated neonatal fMRI using multiband EPI’, in proceedings of international society of magnetic resonance in medicine, p. 3911.

Salimi-Khorshidi, G. et al. (2014) ‘Automatic denoising of functional MRI data: Combining independent component analysis and hierarchical fusion of classifiers’, NeuroImage. Elsevier B.V., 90, pp. 449–468.

Schnabel, J. A. et al. (2001) ‘A generic framework for non-rigid registration based on non-uniform multi-level free-form deformations’, International Conference on Medical Image Computing and Computer-Assisted Intervention, pp. 573–581.

Schuh, A. et al. (2018) ‘Unbiased construction of a temporally consistent morphological atlas of neonatal brain development’, bioRxiv, p. 251512.

Shi, F. et al. (2011) ‘Infant brain atlases from neonates to 1- and 2-year-olds’, PLoS ONE, 6(4), pp. 1–11.

Silbereis, J. C. et al. (2015) ‘The Cellular and Molecular Landscapes of the Developing Human Central Nervous System’, Neuron, 89(2), pp. 248–268.

Smith, R. E. et al. (2012) ‘Anatomically-constrained tractography: Improved diffusion MRI streamlines tractography through effective use of anatomical information’, NeuroImage, 62(3), pp. 1924–1938.

Smith, R. E. et al. (2015) ‘SIFT2: Enabling dense quantitative assessment of brain white matter connectivity using streamlines tractography’, NeuroImage. Academic Press Inc., 119, pp. 338–351.

Sporns, O. (2010) ‘Brain Networks: Structure and Dynamics’, Networks of the Brain.

Sporns, O., Tononi, G. and Kötter, R. (2005) ‘The human connectome: A structural description of the human brain’, PLoS Computational Biology, 1(4), pp. 0245–0251.

Suárez, L. E. et al. (2020) ‘Linking Structure and Function in Macroscale Brain Networks’, Trends in Cognitive Sciences. The Authors, 24(4), pp. 302–315.

Takahashi, E. et al. (2012) ‘Emerging cerebral connectivity in the human fetal Brain: An MR tractography study’, Cerebral Cortex, 22(2), pp. 455–464.

Taoudi-Benchekroun, Y. et al. (2020) ‘Predicting age and clinical risk from the neonatal connectome’, bioRxiv.

Tau, G. Z. and Peterson, B. S. (2010) ‘Normal Development of Brain Circuits’, Neuropsychopharmacology. Nature Publishing Group, 35(10), pp. 147–168.

Tournier, J. D. et al. (2019) ‘MRtrix3: A fast, flexible and open software framework for medical image processing and visualisation’, NeuroImage.

Toyoizumi, T. et al. (2013) ‘A Theory of the Transition to Critical Period Plasticity: Inhibition Selectively Suppresses Spontaneous Activity’, Neuron. Elsevier Inc., 80(1), pp. 51–63.

Tustison, N. J. et al. (2010) ‘N4ITK: Improved N3 bias correction’, IEEE Transactions on Medical Imaging.

Tzourio-Mazoyer, N. et al. (2002) ‘Automated Anatomical Labeling of Activations in SPM Using a Macroscopic Anatomical Parcellation of the MNI MRI Single-Subject Brain’, Neuroimage, 15, pp. 273–289.

Vanderwal, T. et al. (2019) ‘Stability and similarity of the pediatric connectome as developmental outcomes’, bioRxiv, pp. 1–22.

Yeh, F. et al. (2016) ‘Quantifying Differences and Similarities in Whole-Brain White Matter Architecture Using Local Connectome Fingerprints’, PLoS Computational Biology, 12(11), pp. 1–17.

Zhu, K. et al. (2016) ‘Hybrid-Space SENSE Reconstruction for simultaneous Multi-Slice MRI’, IEEE Transactions on Medical Imaging. IEEE, 35(8), pp. 1824–1836.

